# SARS-CoV-2 ORF8 limits expression levels of Spike antigen

**DOI:** 10.1101/2022.11.09.515752

**Authors:** Ik-Jung Kim, Yong-ho Lee, Mir M. Khalid, Yini Zhang, Melanie Ott, Eric Verdin

## Abstract

Survival from COVID-19 depends on the ability of the host to effectively neutralize virions and infected cells, a process largely driven by antibody-mediated immunity. However, with the newly emerging variants that evade Spike-targeting antibodies, re-infections and breakthrough infections are increasingly common. A full characterization of SARS-CoV-2 mechanisms counteracting antibody-mediated immunity is needed. Here, we report that ORF8 is a SARS-CoV-2 factor that controls cellular Spike antigen levels. ORF8 limits the availability of mature Spike by inhibiting host protein synthesis and retaining Spike at the endoplasmic reticulum, reducing cell-surface Spike levels and recognition by anti-SARS-CoV-2 antibodies. With limited Spike availability, ORF8 restricts Spike incorporation during viral assembly, reducing Spike levels in virions. Cell entry of these virions leaves fewer Spike molecules at the cell surface, limiting antibody recognition of infected cells. Our studies propose an ORF8-dependent SARS-CoV-2 strategy that allows immune evasion of infected cells for extended viral production.

## Introduction

Severe acute respiratory syndrome coronavirus 2 (SARS-CoV-2) is the causative agent of COVID-19, a major worldwide pandemic resulting in 6 million confirmed deaths. Several genetic and environmental factors contribute to the survival from COVID-19, with many of them involved in the host capacity to effectively detect and neutralize the virions and the infected cells [1]. Upon entry of SARS-CoV-2 into the host cell, the first line of host defense is innate immunity, sensing the viruses and recruiting immune cells to the initial site of infection in a timely manner [2].

After the first several days in contact with SARS-CoV-2 virions, the immune system develops antibody-mediated humoral immunity, which allows targeted detection of viral antigens on the virions or infected cells [3]. The importance of antibody-mediated immunity against SARS-CoV-2 infection is evident with the high effectiveness of the approved COVID-19 vaccines, which boost production of antibodies against SARS-CoV-2. Specifically, these vaccines were designed to target conserved regions of the Spike protein, a key structural component of SARS-CoV-2 that mediates host cell entry. Upon SARS-CoV-2 infection, a high titer of anti-Spike antibodies develops [4], and the antibody binding to the virions limits the mobility of virions and blocks the host cell entry [5]. These anti-Spike antibodies may also react to Spike molecules on the surface of SARS-CoV-2-infected cells [3], attracting immune cells for phagocytosis or cytotoxicity actions. Targeting both virions and infected cells is important for the maximal antibody activity to antagonize the SARS-CoV-2 dissemination [3].

However, despite a high anti-Spike antibody titer in COVID-19 convalescent or vaccinated individuals, infections in these individuals are increasingly becoming common, suggesting the possibility that several SARS-CoV-2 mechanisms exist to manipulate or evade antibody-mediated immunity. In support of this idea, the superior fitness of new variants of concern (VOCs) that are now dominant worldwide largely derives from mutations on Spike that limits antibody affinity [6]. To respond effectively to the continued emergence of increasingly evasive VOCs, further investigations are required to fully characterize the SARS-CoV-2 mechanisms for limiting antibody-mediated immunity.

Here, we report that ORF8, a SARS-CoV-2 protein that is largely uncharacterized, has a potential pro-viral role by controlling the availability of Spike antigens during infection. We found that ORF8 is a luminal protein of the endoplasmic reticulum (ER) that strongly interacts with Spike. With ORF8, Spike protein levels were diminished (similarly by the VOC genotype ORF8 S84L)) by two independent mechanisms: 1) ORF8 limits the host capacity to synthesize proteins, and 2) covalent interactions with Spike inhibit translocation of Spike to the Golgi. With the limited availability of mature Spike, ORF8 also limited the abundance of cell-surface Spike, a trigger for fragment crystallization (Fc) receptor functions that can be initiated by anti-SARS-CoV-2 human sera. Viral particles produced in cells co-expressing ORF8 incorporate less Spike and exhibit lower infectivity. However, infection with these viral particles results in much lower levels of virus-derived Spike molecules at the cell surface, limiting the reactivity of the anti-SARS-CoV-2 human sera. Our studies provide evidence that supports the model that ORF8 contributes to extended viral production by tightly controlling the availability of Spike antigens in infected cells or virions, evading immune detection of infected cells.

## Results

### SARS-CoV-2 ORF8 is an ER luminal protein

ORF8 interacts with an array of ER chaperone proteins [7], suggesting that ORF8 is subcellularly localized to the ER. Computational analysis (Protter) of the amino acid sequence of ORF8 predicts that the first 16 N-terminal amino acids are an ER signal peptide (Fig 1A), suggesting that, upon *de novo* synthesis, ORF8 is translocated into the ER. To test this possibility, A549, a human lung epithelia-derived cell line, transfected with a plasmid encoding C-terminal double Strep-tagged ORF8 (ORF8-Strep), was fixed, permeabilized and immunostained for Strep, and disulfide isomerase (PDI) (ER-specific organelle marker). ORF8 (green signals) visually colocalized with protein PDI (red signals) (Fig 1B), as manifested by a high degree of similarity between the two signal intensities (Fig 1C) along the cross-sectional arrow (Fig 1B). The possibility that ORF8 is an ER protein was further evaluated by biochemical studies. HEK293T cells transfected with a plasmid encoding C-terminal Flag-tagged ORF8 (ORF8-Flag) were subcellularly fractionated by differential centrifugation, yielding major cellular compartment fractions (e.g., ER, mitochondria, and cytosol) (Fig 1D), and those fractions were evaluated by immunoblot analyses for ORF8-Flag signals. The ORF8 signal was observed only in the ER fractions (characterized by Calnexin), but not in mitochondria (COX4) or cytosol (β-actin), indicating that ORF8 is predominantly localized to ER within cells.

**Fig 1.**
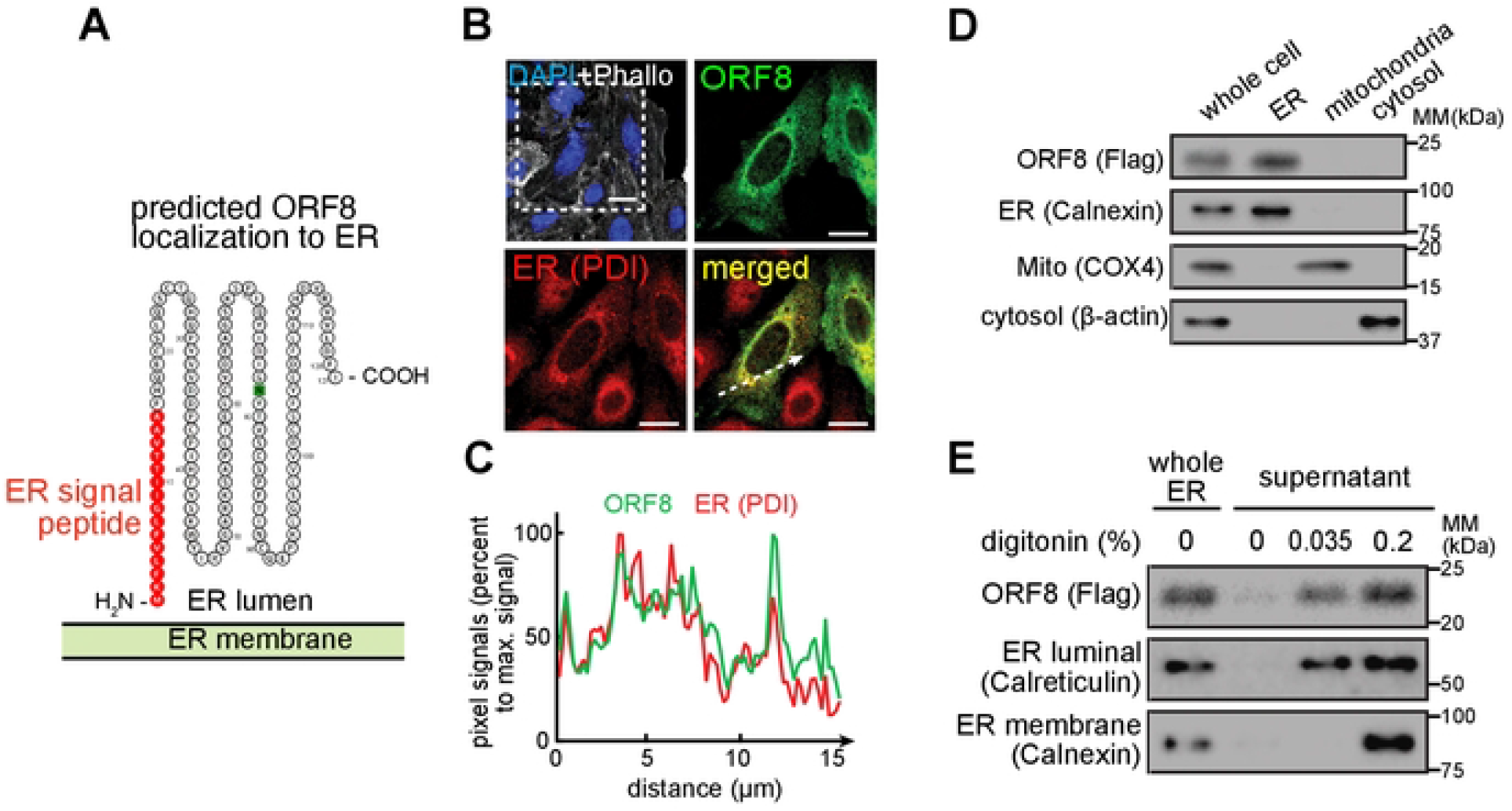
SARS-CoV-2 ORF8 is an ER luminal protein. **A.** Amino acid sequence analysis (Protter) of ORF8 and prediction as an ER luminal protein (due to the presence of ER signal peptide and the absence of a transmembrane domain). **B.** A549 cells transfected with a plasmid encoding ORF8-Strep were fixed, permeabilized, and immunostained for Strep (ORF8) or PDI (ER marker), which were analyzed by fluorescence confocal microscopy imaging. The cells were counterstained using DAPI and Phalloidin (upper left). The dashed box is digitally enlarged to show colocalization (bottom right) of ORF8 (upper right) and ER (bottom left). White scale bars = 10 μm. **C.** The pixel intensities of ORF8 and PDI along the dashed arrow in **B** are plotted. **D.** HEK293T cells transfected with a plasmid encoding ORF8-Flag were mechanically lysed and fractionated by differential centrifugation. The indicated subcellular fractions were evaluated by immunoblot analysis for Flag (ORF8), Calnexin (ER marker), COX4 (mitochondrial marker) or ß-actin (cytosolic marker). **E.** The ER subcellular fractions in **D** were further incubated with two concentrations of digitonin (0, 0.035, or 0.2%). The fractions were then centrifuged, and the supernatants containing digitonin-solubilized proteins were evaluated by immunoblot analysis as in **D**. **B–E.** The data represent three independent experiments.

Lacking a transmembrane domain (Fig 1A), we predicted that ORF8 is a luminal protein after translocating to the ER. To test the prediction, the ORF8-containing ER fractions collected previously (Fig 1D) were incubated with two concentrations of digitonin. With the lower concentration (0.035%), Calreticulin (ER luminal marker) was solubilized and remained in the supernatant after high-speed centrifugation. With the higher concentration (0.2%), both Calreticulin and Calnexin (ER membrane marker) remained in the supernatant (Fig 1E). After a 45-min incubation with the indicated concentrations of digitonin, the fractions were centrifuged, and the proteins in the supernatant were examined by immunoblot analysis. The ORF8-Flag signals were noted at the lower digitonin concentration (0.035%), consistent with the hypothesis that ORF8 is an ER luminal protein.

### ORF8 modulates Spike protein levels

Three SARS-CoV-2 proteins (i.e., Spike, ORF7a, ORF8) contain an ER signal peptide, and Spike is a key viral component highly implicated in the viral infectivity. With ORF8 and Spike existing in the same subcellular space of the ER, as manifested by the colocalization of ORF8 and Spike signals (Fig 2A), we investigated the possibility that ORF8 alters Spike levels. HEK293T cells co-transfected with plasmids encoding C-terminal Flag-tagged Spike (Spike-Flag), and ORF8-Strep or eGFP-Strep (negative control) were lysed for immunoblot analyses (Fig 2B and 2C) with an antibody targeting the Spike S2 or S1 region (Fig 2D). Two immunoblot bands were detected for Spike (Fig 2B), corresponding to uncleaved nascent Spike (220 kDa), and the Spike that is cleaved (90 kDa in αS2 blot, and 130 kDa in αS1 blot) at the furin-cleavage site, a reaction thought to occur at the ER-Golgi intermediate complex (ERGIC) or Golgi [8], resulting in S1 and S2 fragments (Fig 2B and 2D). The total Spike levels (calculated by combining uncleaved and S2 signals) decreased (> 50%) in an ORF8-dependent manner (Fig 2C). Moreover, the band intensities corresponding to S2 or S1 fragments decreased to a greater extent (> 95% decrease) in an ORF8-dependent manner (Fig 2C). No other immunoblot bands were detected under our experimental conditions (Fig S1A), validating our quantitative measurement of Spike protein levels. The ORF8-dependent modification of Spike protein levels was reproduced using non-tagged Spike and ORF8 (Fig S1B), validating the use of the C-terminal tagged constructs for our investigation.

**Fig 2.**
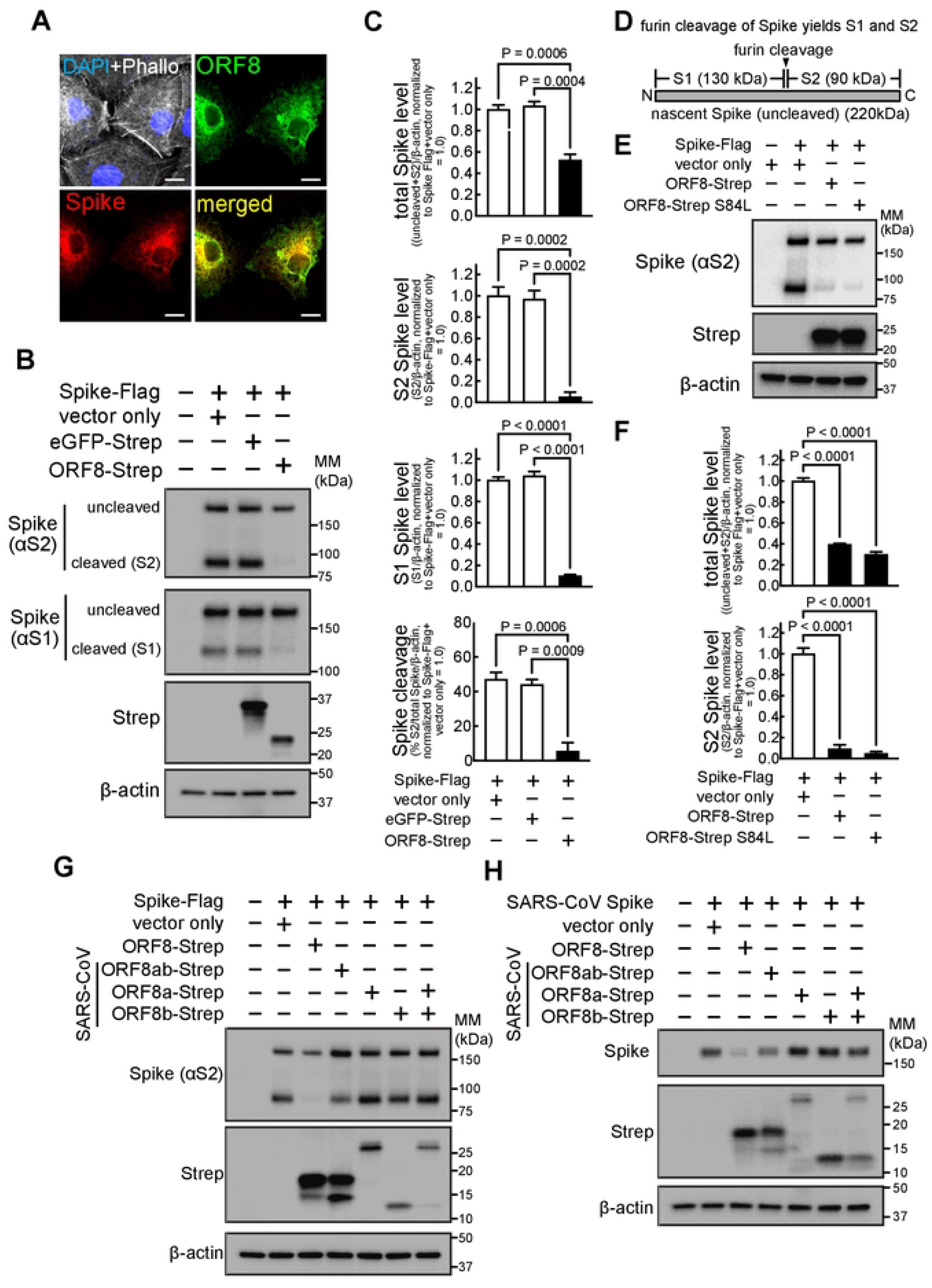
ORF8 colocalizes with Spike and modulates Spike protein levels and furin-dependent processing. **A.** A549 cells co-transfected with plasmids encoding Spike-Flag or ORF8-Strep were fixed, permeabilized, and immunostained for Flag (Spike) and Strep (ORF8), which were analyzed by fluorescence confocal micros∞py imaging. The cells were counterstained using DAPI and Phalloidin, (upper left), with ORF8 (upper right), Spike (bottom left), or both ORF8 and Spike signals merged (bottom right). White scale bars = 10 μm. **B**, **C**, **E–H.** HEK293T cells co-transfected with plasmids encoding Spike-Flag (**B**, **C**, **E–G**) or SARS-CoV-derived Spike (**H**), or, eGFP-Strep (**B**, **C**), ORF8-Strep (**B**, **C**, **E–H**), ORF8-Strep S84L (the B lineage genotype) (**E** and **F**), or SARS-CoV-derived ORF8-Strep genotypes (ORF8ab, ORF8a, ORF8b, or ORF8a and ORF8b together) (**G** and **H**) were lysed for immunoblot analysis using antibodies against S2 (detects uncleaved and S2 fragment of Spike, or SARS-CoV Spike) (**B**, **C**, **E-H**), S1 fragment (detects uncleaved and S1 fragment of Spike) (**B**, **C**), Strep (detects eGFP or ORF8s) (**B**, **C**, **E–H**), and ß-actin (**B**, **C**, **E–H**). **D.** Depiction of whole Spike with the site that can be cleaved by furin, yielding S1 and S2 fragments. Apparent immunoblot mass (kDa) is indicated. The data represent or are combined from three independent experiments and are presented as mean ± s.d. Statistical significance was analyzed by one-way ANOVA (Dunnett’s test).

Next, we determined if our findings could be extended to the recent emergence of VOCs. The amino acid sequence of ORF8 is highly conserved across different sub-strains, except for the S84L mutation [9] that is commonly found in the major VOCs, including Delta and Omicron sub-variants. ORF8-Strep with S84L mutation (ORF8-Strep S84L) also altered Spike protein levels similarly (Fig 2E and 2F), suggesting that the ORF8 actions on Spike are conserved in the VOC ORF8 genotypes. Finally, the ORF8 alternation of Spike protein levels was not observed with the ORF8s derived from SARS-CoV (ORF8ab) (Fig 2G) or ORF8a and ORF8b, which later emerged by truncation of 29 amino acids [10], and minor reduction by ORF8ab when paired with their own SARS-CoV Spike (Fig D). These findings suggest that the ORF8 modulation of cellular Spike levels is a SARS-CoV-2-specific mechanism.

### ORF8 covalently interacts with Spike at the ER and impedes Spike translocation to the Golgi

Next, we investigated whether ORF8 and Spike in the ER interact by creating an ORF8-Flag construct with an I9P mutation (ORF-Flag I9P) that disrupts the α-helix structure of the ER signal peptide by introducing a proline kink. Loss of ability to translocate to the ER was validated by immunofluorescence microscopy analysis, as manifested by the cytosolic distribution of ORF8-Flag I9P or ORF8-Flag lacking the entire ER signal peptide (ORF8-Flag Δ1-17) (green signals) (Fig S2A), as well as the loss of ER (red signals) colocalization with ORF8-Flag I9P or ORF8-Flag Δ1-17 (Fig S2A). Furthermore, immunoblot analysis under non-reducing conditions (to preserve disulfide bonds) showed the non-mutated ORF8-Flag as multiple bands (Fig S2B), which is attributed to intermolecular disulfide bonds that form within the oxidizing ER lumen environment [11], whereas ORF8-Flag I9P and ORF8-Flag Δ1-17 were observed as a single band.

To evaluate the importance of the ORF8 localization to the ER for its effect on Spike protein levels, cells were co-transfected with a plasmid encoding non-tagged Spike, and a plasmid encoding GFP-Flag (negative control), ORF8-Flag, or ORF-Flag I9P. The cells were lysed, the lysates were incubated with anti-Flag magnetic beads, and the immunoprecipitated proteins were analyzed by western blotting. We observed a loss of cleaved S2 fragment of Spike in cells co-expressing ORF8-Flag (Fig 3A, input), but not in cells co-expressing GFP-Flag control or ORF8-Flag I9P. Moreover, Spike was detected in the immunoprecipitated samples collected from cells co-expressing ORF8-Flag, but nor GFP-Flag control or ORF8-Flag I9P (Fig 3A), indicating that Spike co-immunoprecipitated with ORF8-Flag but not with ORF-Flag I9P. These studies support the model that ORF8 interacts with Spike at the ER, and that ORF8 translocation to the ER is required for the ORF8-Spike interaction and for altering Spike protein levels.

**Fig 3.**
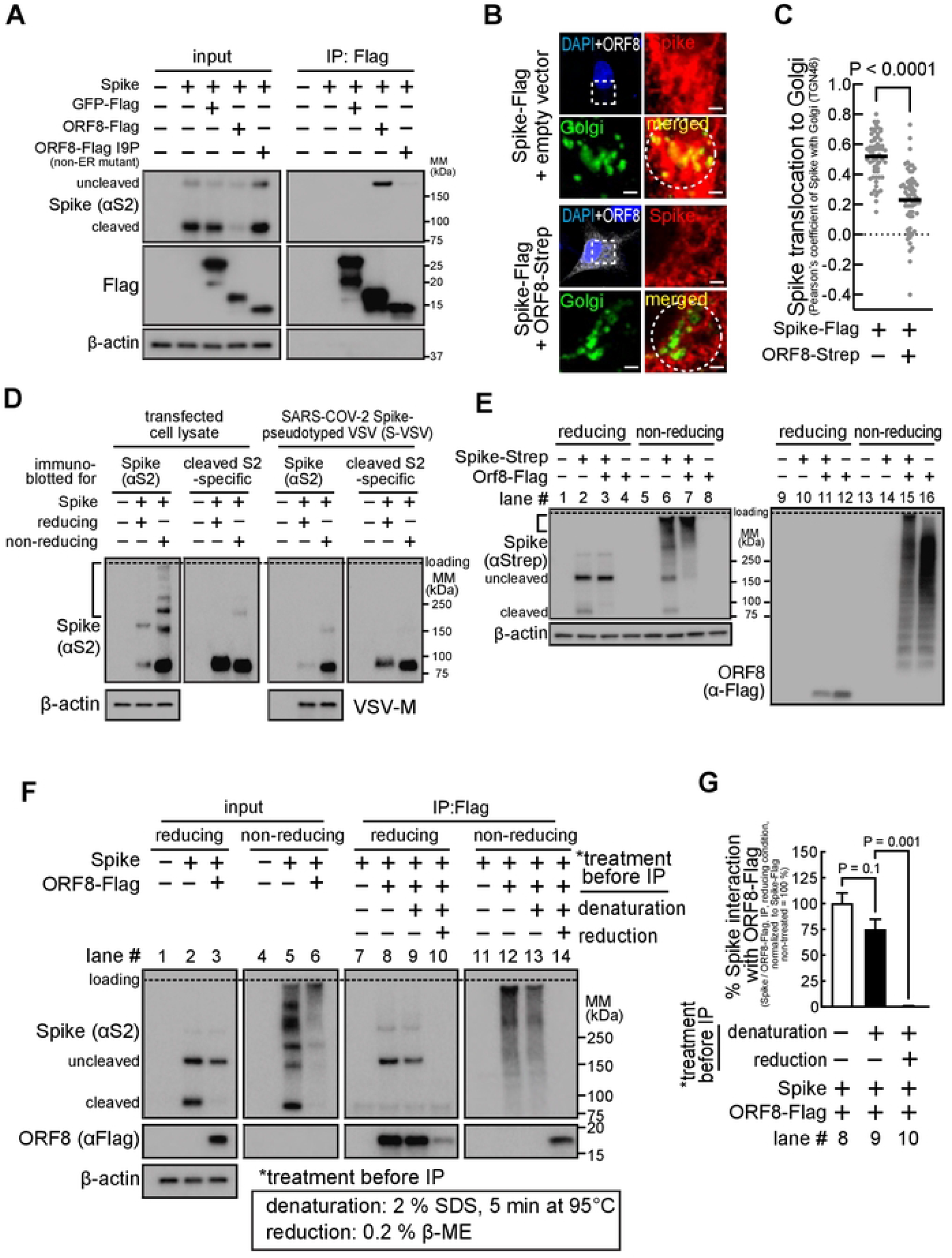
ORF8 covalently interacts with Spike and hampers Spike translocation to the Golgi apparatus. **A**, **D–G**. HEK293T cells co-transfected with plasmids encoding non-tagged Spike (**A**, **D**, **F**, **G**), or Spike-Strep (**E**), and/or GFP-Flag (**A**), ORF8-Flag (**A**, **E**, **F**, **G**), ORF8-Flag I9P (non-ER mutant) (**A**), or ORF8-Flag Δ1-17 (ER signal deletion) (**A**1) were not infected (**A**, **D–G**) or infected (**D**) with VSVΔ G-GFP, for production of a hybrid VSVΔG-GFP that incorporated fully mature SARS-CoV-2 Spike (S-VSV). The cells above expressing Spike and/or ORF8 constructs, or, S-VSV collected from the culture medium were lysed and directly analyzed by immunoblots using antibodies against S2 (detects uncleaved and S2 fragment of Spike) (**A**, **D**, **F**), N-terminus of S2 (detects S2 fragment only) (**A**, **D**, **F**), Flag (detects GFP or ORF8) (**A**, **E**, **F**, **G**), Strep (detects Spike) (**E**), and ß-actin, under reducing or non-reducing (protein interactions through disulfide bonds were preserved) conditions or further incubated with anti-Flag magnetic beads (**A**, **F**, and **G**), without (**A**, **F**, and **G**) or with pre-treatment (**F** and **G**) (denaturation; 2% SDS, 5 min at 95 °C, reduction: 0.02% ß-ME) of the cell lysates. The proteins that were immunoprecipitated were analyzed by immunoblots under reducing (**A**, **F**, **G**) or non-reducing condition (**F** and **G**). **B**, **C**. A549 (**B**) or HEK293T (**C**) cells co-transfected with a plasmid encoding Spike-Flag and a bicistronic plasmid encoding ORF8 and eGFP (ORF8-Strep-IRES-eGFP) were fixed, permeabilized, and immunostained using antibodies against S2 (detects uncleaved and S2 fragment of Spike) and TGN46 (Golgi marker), which were analyzed by fluorescence confocal microscopy imaging. **B**. The cells that were not expressing or expressing ORF8-Strep (identified by eGFP signals, pseudo-∞lored to white) were counterstained using DAPI (upper left), with the dashed box that is digitally enlarged to evaluate colocalization (bottom right) of Spike (upper right) and Golgi (TGN46) (bottom left). White scale bars = 2 μm. **C**. Colocalization of Spike and Golgi signals within the circular area encompassing Golgi (dashed circles in **B**) was analyzed by measuring Pearson’s coefficient in 60 cells (combined from three independent experiments, in which 20 cells were randomly selected). The data represent or are combined from three independent experiments and are presented as mean ± s.d. Statistical significance was analyzed using two-tailed Student’s t test (**C**) or one-way ANOVA (Dunnett’s test) (**G**).

More cleaved Spike-Flag was lost (> 95%) than total Spike (> 50%) (Fig 2B and 2C), and the Spike cleavage rate was lower (Fig 2B and 2C) in cells co-expressing ORF8-Strep, suggesting that furin cleavage of Spike is inhibited by ORF8. The furin-dependent Spike cleavage (Fig 2D) is a post-ER event that occurs at the ERGIC or Golgi. Thus, we hypothesized that ORF8 interaction with Spike at the ER inhibits Spike translocation to Golgi, preventing furin-cleavage. In support of this model, the Spike species that interacts with ORF8 is uncleaved (Fig 3A). To further investigate whether Spike translocation to the Golgi is altered by ORF8, A549 or HEK293T cells co-transfected with plasmids encoding Spike-Flag and a bicistronic plasmid encoding both ORF8-Strep and eGFP separated by internal ribosomal entry cite (IRES) (ORF8-Strep-_IRES_-eGFP) were fixed, permeabilized, immunostained for Spike S2 and trans-Golgi network protein 46 (TGN46) (used as a Golgi marker), and examined by confocal microscopy. Spike (red signals) colocalization to the Golgi (green signals) decreased visually (Fig 3B) and quantitatively (Fig 3C) (calculated by Pearson’s coefficient) in cells co-expressing ORF8-Strep (detectable by eGFP signal (pseudo-colored to white)). These studies support the model that Spike interaction with ORF8 retains itself at the ER and impedes its translocation to Golgi.

Interestingly, Spike protein expression is largely detected as high-molecular-mass smear under non-reducing conditions (bracket, Fig 3D). This was not the case under reducing conditions (Fig 3D) (suggesting the smear is Spike species aggregated through disulfide bonds), or when immunoblotted with an antibody that detects cleaved S2 Spike only (Fig 3D) (suggesting the smear is uncleaved Spike), or with the fully mature Spike molecules incorporated into viral particles (Fig 3D) (suggesting the smear is Spike still undergoing maturation). These observations suggest that the smear represents the uncleaved Spike molecules undergoing protein folding at the ER. We hypothesized that Spike retention at the ER (Fig 3B and 3C) within ORF8-coexpressing cells resulted from interaction with the cysteine-rich ORF8 (5.8%,7/121 residues), and we first tested whether ORF8-Spike interaction involves covalent bonds. Cells co-transfected with plasmids encoding ORF8-Flag and Spike-Strep were lysed and evaluated by immunoblot under non-reducing conditions for the molecular mass distribution of ORF8-Spike complexes (Fig 3E). Both Spike-Strep (lane #: 6) and ORF8-Flag (lane #: 16) signals were generally upshifted towards the higher molecular mass species (bracket, lane #: 7, 15) than cells singly expressing Spike-Strep only or ORF8-Flag only, indicating formation of higher molecular mass, disulfide bond–based protein aggregates. Notably, the two non-intermolecular Spike-Strep bands (cleaved/uncleaved, lane #: 6) in cells singly expressing Spike-Strep were barely detected in cells co-expressing Spike-Strep and ORF8-Flag (lane #: 7), suggesting that most cellular Spike molecules remain aggregated through disulfide bonds in cells co-expressing ORF8.

To directly test whether the ORF8-Spike interaction is mostly associated with disulfide bonds, cells co-transfected with plasmids encoding Spike or ORF8-Flag were lysed and the cell lysates were pre-incubated at 95°C for 5 min in 2% SDS (to break up non-covalent protein-protein interactions) and in the absence or presence of 0.2% β-ME (to break up intra- and inter-molecular disulfide bonds). After the pre-incubation, the lysates were immunoprecipitated with anti-Flag magnetic beads and analyzed by immunoblotting under non-reducing or reducing conditions (Fig 3F and 3G). We observed co-immunoprecipitation of Spike even after the denaturation (lane # 9), at a level that is not significantly different from the same lysates that were not pre-incubated (lane #: 8). The Spike co-immunoprecipitation was completely abolished under reducing conditions (lane #: 10), suggesting that ORF8-Spike interaction is predominantly established through disulfide bonds. Furthermore, the co-immunoprecipitated Spike under non-reducing conditions was entirely detected as high-molecular-mass smears (lane #: 12) that were retained even under denaturing conditions (lane #: 13). These studies support the model that Spike and ORF8 form protein aggregates through disulfide bonds at the ER, and Spike translocation to Golgi is impeded.

### Host protein synthesis is inhibited within cells expressing ORF8

We further tested this proposed mechanism with decanoyl-RVKR-CMK (or simply CMK), a furin inhibitor (Fig 4A). However, the decrease in total Spike-Flag levels (uncleaved + cleaved S2) in cells co-expressing ORF8-Strep was not clearly manifested in cells incubated with CMK. Moreover, total levels of a modified Spike-Flag insensitive to furin cleavage (the furin cleavage site was deleted) (Spike-Flag FKO) [12] decreased similarly (> 50%) in cells co-expressing ORF8-Strep (Fig 4B and 4C), suggesting an additional ORF8 mechanism responsible for the total Spike decrease. We first evaluated whether Spike expression is modulated by ORF8 at the transcription levels, but no decrease in the transcript levels of Spike-Flag was detected by RT-qPCR in cells co-expressing ORF8-Strep (Fig 4D). Interestingly, flow cytometry analysis of cells transfected with a bicistronic plasmid encoding ORF8-Strep-_IRES_-eGFP showed significantly lower eGFP expression (Fig 4E). ORF8 inhibition of eGFP expression suggested the possibility that ORF8 might limit the host capacity for protein synthesis. To investigate this possibility, cells transfected with a bicistronic plasmid encoding no ORF8 (empty-_IRES_-eGFP) or ORF8-Strep genotypes (ORF8-Strep-_IRES_-eGFP or ORF8-Strep S84L-_IRES_-eGFP) were incubated with L-homopropargylglycine (HPG), a Click-modified methionine analog that is incorporated into newly synthesized proteins. After 30 min, cells were harvested, fixed, and permeabilized, and the incorporated cellular HPG was fluorescently labeled for detection by flow cytometry. HPG incorporation (< 15%) in cells expressing ORF8-Strep or ORF8-Strep S84L (eGFP-positive) was much less than in cells not expressing ORF8 (eGFP-positive) (Fig 4F and 4G), supporting the hypothesis that ORF8 inhibits global host protein synthesis. Lastly, no significant reduction in HPG incorporation was observed with SARS-CoV ORF8a-Strep or ORF8b-Strep, and only a minor reduction in cells expressing ORF8ab-Strep (33 %) (Fig 4H), suggesting the ORF8-dependent host protein synthesis inhibition is a unique feature of SARS-CoV-2.

**Fig 4.**
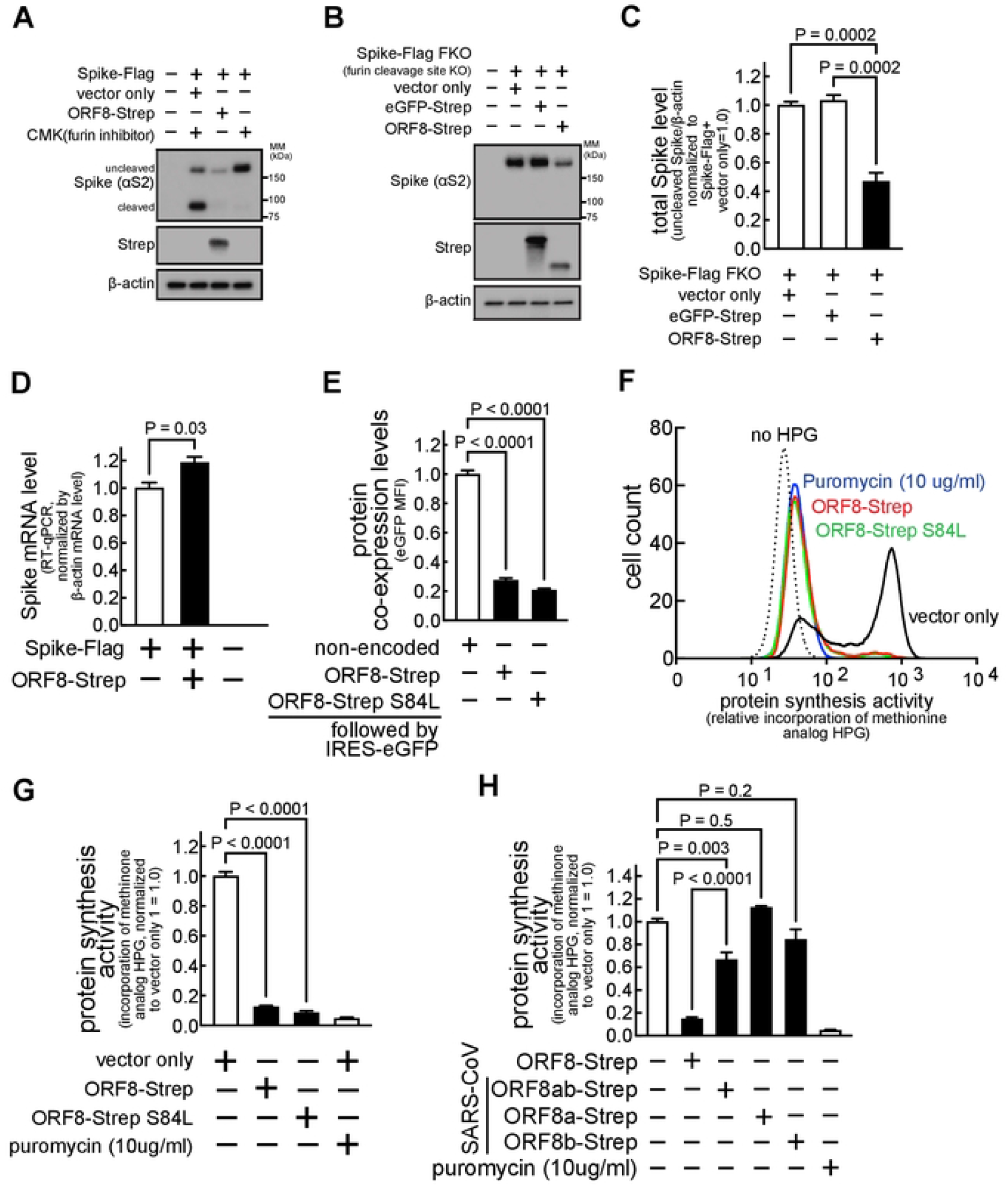
Host protein synthesis is inhibited in cells expressing ORF8. **A–D**. HEK293T cells co-transfected with a plasmid encoding Spike-Flag (**A** and **D**) or Spike-Flag FKO (furin cleavage site KO) (**B** and **C**), and eGFP-Strep (**B** and **C**) or ORF8-Strep (**A–D**), were lysed after incubation without (**B–D**) or with (**A**) CMK (furin inhibitor), and evaluated by immunoblot analysis (**A–C**) using antibodies against S2 (detects uncleaved and S2 fragment of Spike), Strep (detects GFP or ORF8), or ß-actin, or, evaluated by RT-qPCR (D) using primers that are designed against Spike. **E–H**. HEK293T cells transfected with a bicistronic plasmid encoding both eGFP and different ORF8-Strep genotypes (ORF8-Strep, ORF8-Strep S84L, or, SARS-CoV-derived ORF8ab-Strep, ORF8a-Strep, or ORF8b-Strep) were incubated in the absence (**E**) or presence (**F–H**) of HPG (methionine analog) without or with puromycin (protein synthesis inhibitor) (**F–H**). After 30 min, the cells were harvested, fixed, and directly analyzed by flow cytometry for the fluorescence signals of eGFP (**E**) or permeabilized after fixation and fluorescently labeled for flow cytometry analysis of the incorporated cellular HPG within cells expressing ORF8-Strep (eGFP-positive) (**F–H**). The data represent or are combined from three independent experiments and are presented as mean ± s.d. Statistical significance was analyzed using one-way ANOVA (**C**, **E**, and **G**; Dunnett’s test, **H**; Tukey’s test) or two-tailed Student’s t test (**D**).

### ORF8 limits cell-surface Spike levels

Once Spike molecules arrive at Golgi after full maturation (as the cleaved form), they are utilized for viral assembly (Fig 3D) or translocated to the host cell surface [13]. With our previous finding that cellular levels of mature Spike decrease in an ORF8-dependent manner (Fig 2B and 2C), we hypothesized that ORF8 might decrease Spike abundance at the cell surface. We first evaluated syncytia (cell-cell fusion) formation, which occurs during SARS-CoV-2 infection [14] by interaction of cell-surface Spike with the ACE2 receptors in neighboring cells. HEK293T cells that stably express ACE2 and TMPRSS2 (HEK293T A/T) [15] were co-transfected with plasmids encoding Spike-Flag or ORF8-Strep. After 18 h, cells were fixed, permeabilized, and immunostained for Flag (Spike) and Strep (ORF8) (Fig 5A). Clear syncytia were formed in cells expressing Spike-Flag, as manifested by collapsed cellular boundaries and multinuclear arrangement (inset). In contrast, cells co-expressing Spike-Flag and ORF8-Strep remained well separated (inset). The inhibition of syncytia formation in cells co-expressing ORF8 suggests reduction in cell-surface Spike levels.

**Fig 5.**
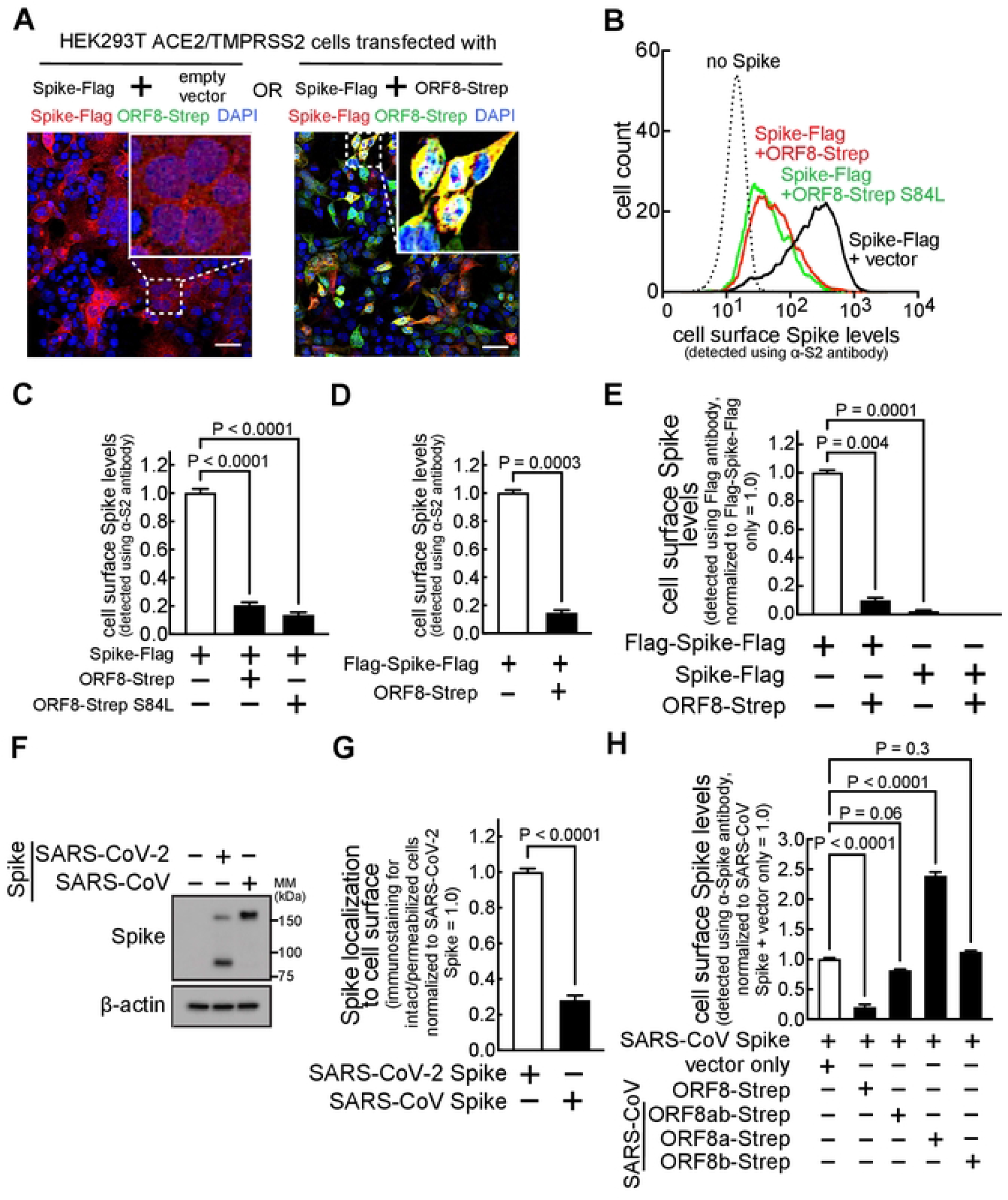
ORF8 limits the levels of cell-surface Spike. **A**. A monolayer of HEK293T cells stably expressing ACE2/TMPRSS2 were co-transfected with a plasmid encoding Spike-Flag or ORF8-Strep. The cells were fixed, permeabilized, and immunostained with antibodies against Flag (Spike) and Strep (ORF8). After counterstaining with DAPI, the cells were analyzed using fluorescence confocal microscopy imaging. The dotted boxes were digitally enlarged (upper right inset). The scale bars = 50 μm. **B–H**. HEK293T cells co-transfected with a plasmid encoding Spike-Flag (C-terminal tagged) (**B**, **C**, **E**), Flag-Spike-Flag (N- and C-terminal tagged) (**D** and **E**), non-tagged Spike (**F** and **G**), or SARS-CoV-derived Spike (**F–H**), and a bicistronic plasmid en∞ding eGFP and different ORF8-Strep genotypes (ORF8-Strep (**B–E**, **H**), ORF8-Strep S84L (B and C) or SARS-CoV-derived ORF8ab-Strep, ORF8a-Strep, or ORF8b-Strep (**H**)). The cells were harvested and directly immunostained for cell surface Spike by incubating with antibodies against S2 (**B–D**, **G** and **H**; detects uncleaved Spike, S2 fragment, or SARS-CoV-derived Spike) or Flag (**E**) or fixed, permeabilized and immunostained by incubating with antibodies against S2 (**G**). Cell-surface Spike levels (B-H) in viable (LIVE/DEAD-negative) cells expressing ORF8-Strep (eGFP-positive) or total cellular Spike levels (**G**) in cells expressing ORF8-Strep (eGFP-positive) were measured by flow cytometry. Cellular expression levels of SARS-CoV-2 Spike or SARS-CoV Spike were confirmed by immunoblot analysis (**F**). The data represent or are combined from three independent experiments and presented as mean ± s.d. Statistical significance was analyzed using one-way ANOVA (**C**, **E** and **H**; Dunnetfs test) or two-tailed Student’s t test (**D** and **G**).

To directly evaluate cell-surface Spike levels, HEK293T cells co-transfected with a plasmid encoding Spike-Flag and a bicistronic plasmid encoding ORF8-Strep genotypes (ORF8-Strep-_IRES_-eGFP, ORF8-Strep S84L-_IRES_-eGFP) were harvested and immunostained using an antibody against Spike S2, followed by incubation with a fluorophore-conjugated secondary antibody as well as a LIVE/DEAD cell viability dye that selectively stains non-viable cells. The viable (LIVE/DEAD-negative) and transfection-positive cells (eGFP-positive) that express no ORF8 or ORF8-Strep, were evaluated by flow cytometry for the abundance of cell surface Spike. Cell-surface Spike signals were greatly reduced (> 80%) in cells co-expressing ORF8-Strep or ORF8-Strep S84L, compared to cells co-expressing no ORF8-Strep (Fig 5B and 5C).

These findings were further validated using a N- and C-terminal-tagged Spike construct (Flag-Spike-Flag) [12], which similarly decreased at the cell surface in an ORF8-dependent manner (Fig 5D). The same experiment, using Flag-Spike-Flag and an anti-Flag antibody that has no access to the cytosolic C-terminal Flag of cell-surface Spike in viable cells, showed no significant signals in cells expressing Spike-Flag, compared to cells expressing Flag-Spike-Flag (Fig 5E). These results indicated that our signal detection is specific to cell-surface-exposed Spike, validating our measurement of cell-surface Spike levels. Moreover, the Flag signals in cells expressing Flag-Spike-Flag, which was thereby corresponding to the N-terminal Flag of cell-surface Spike, significantly decreased (> 90%) by ORF8 co-expression (Fig 5E). These studies demonstrated that levels of N-terminal S1 fragment of cell-surface Spike also decreases in an ORF8-dependent manner.

Lastly, the SARS-CoV Spike, which was expressed to a similar level as SARS-CoV-2 Spike (Fig 5F), was detected in much lower levels at the cell surface (normalized by the total Spike levels) (Fig 5G), and no significant reduction of cell-surface SARS-CoV Spike levels was detected in cells co-expressing the SARS-CoV-derived ORF8 genotypes (Fig 5H), demonstrating that reduction of cell-surface Spike levels is a SARS-CoV-2 ORF8-specific phenomenon.

### ORF8 limits the reactivity of anti-SARS-CoV-2 human sera towards Spike-producing cells

To understand the biological consequence of altered cell-surface Spike levels, we determined if the ORF8 reduction of cell-surface Spike levels interferes with antibody-mediated immune detection of infected cells, a reaction triggered by binding of humoral anti-SARS-CoV-2 antibodies to cell-surface antigens. We next sought to evaluate ORF8’s effect on the ability of anti-SARS-CoV-2 human sera to trigger Fc receptor functions. Cells were co-transfected with a plasmid encoding Spike and a bicistronic plasmid encoding ORF8-Strep-_IRES_-eGFP and harvested and incubated with sera collected from three COVID-19 convalescent (Fig 6A and 6B) or three COVID-19 negative (Fig 6A) human donors. This was followed by incubation with a LIVE/DEAD cell viability dye and a fluorophore-conjugated secondary antibody that specifically detects the Fc region of human immunoglobulin G (IgG) molecules. Flow cytometry showed strong reactivity of the convalescent sera towards Spike-expressing cells: human IgG Fc signals were up to 55-fold greater in cells (LIVE/DEAD-negative, eGFP-positive) incubated with the convalescent sera (Fig 6A) than cells incubated with the COVID-19 negative sera. The signals were significantly lower in cells co-expressing Spike-Flag and ORF8-Strep (< 80 %) (Fig 6B), supporting the model that the reactivity of the convalescent sera to the cell-surface Spike is limited by ORF8.

**Fig 6.**
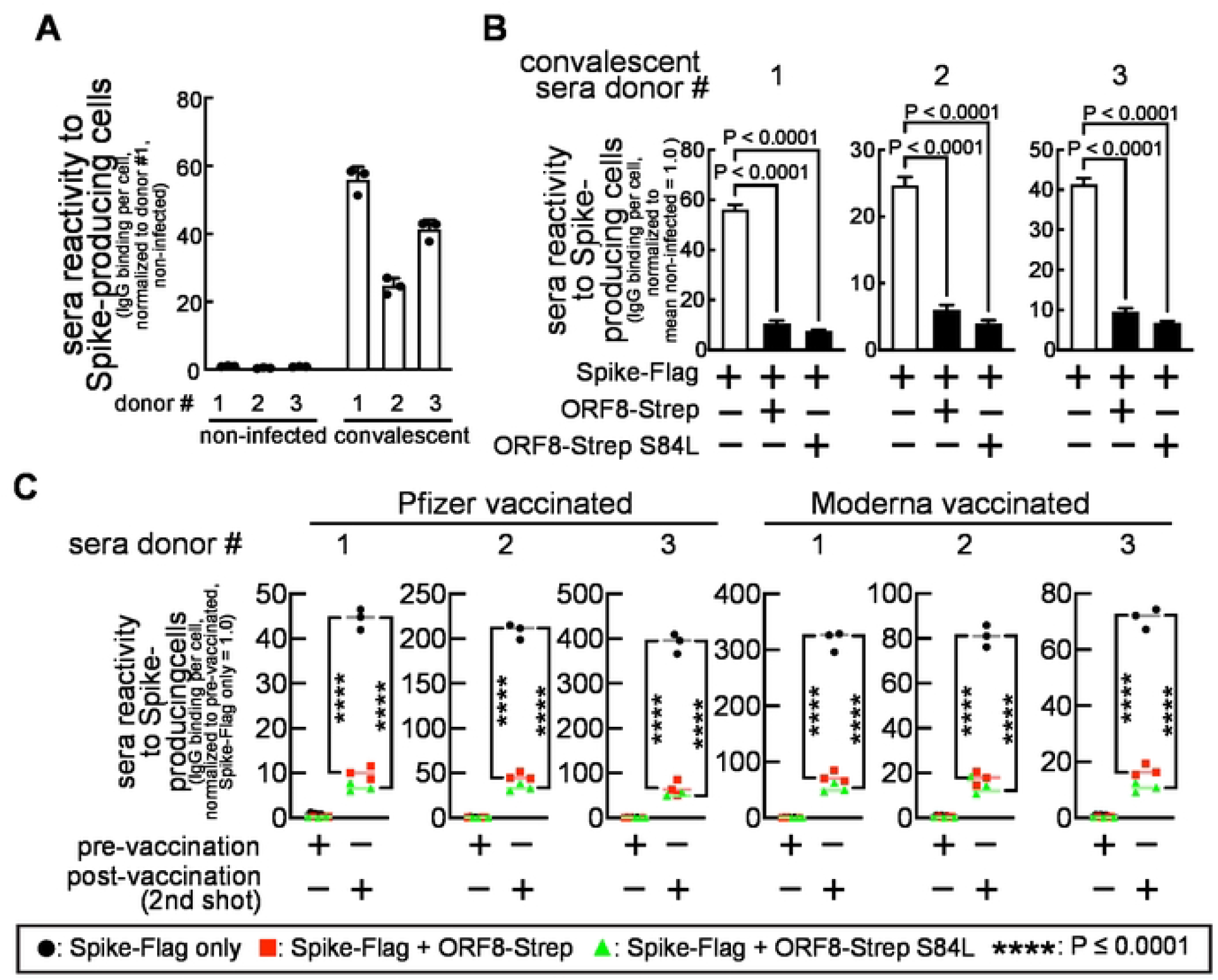
ORF8 limits the reactivity of anti-SARS-CoV-2 (convalescent or vaccinated) human sera towards Spike-producing cells. **A–C**. HEK293T cells ∞-transfected with a plasmid encoding Spike-Flag and a bicistronic plasmid en∞ding mCherry and ORF8-Strep genotypes (ORF8-Strep, ORF8-Strep S84L) were harvested and incubated with anti-SARS-CoV-2 human sera collected from three COVID-19 convalescent (**A**, **B**), six vaccinated (**C**) (three Pfizer and three Moderna, before 1st shot (pre-vaccination) and after 2nd shot (post-vaccination)), or three COVID-19-negative donors (**A**). The IgG molecules in the sera that reacted to Spike-producing cells were fluorescently labeled using antibodies that specifically recognize the Fc region of human IgG molecules. The human IgG Fc levels in viable (LIVE/DEAD-negative) cells expressing ORF8-Strep (mCherry-positive) were evaluated by flow cytometry. The data are combined from three independent experiments and represented as mean ± s.d. Statistical significance was analyzed using one-way (**B**, Dunnett’s test) or two-way AN OVA (**C**, Tukey’s test).

Next, we determined if our findings can be extended to vaccinated individuals. The same experiment was completed with sera from six vaccinated (three Pfizer- and three Moderna-vaccinated, pre-vaccination (collected before the 1^st^ shot) and post-vaccination (collected after the 2^nd^ shot)) human donors. Human IgG Fc signals in cells incubated with the post-vaccination sera were dramatically greater (up to 400-fold) than cells incubated with pre-vaccination sera, regardless of the vaccine brands (Fig 6C), and the signals were decreased in cells co-expressing Spike-Flag and ORF-Strep (> 80%) (Fig 6C). These results indicate that the anti-SARS-CoV-2 human sera, both convalescent and vaccinated, reacts less to the cells co-expressing Spike and ORF8, and their capacity to trigger Fc receptor functions is limited, supporting the model that ORF8 contributes to the survival of SARS-CoV-2-infected cells from the antibody-mediated immunity.

### ORF8 restricts Spike incorporation during viral assembly and reduces viral infectivity, but limits the reactivity of anti-SARS-CoV-2 human sera towards the infected cells

Next, we examined the effect of ORF8 on mature Spike molecules utilized for viral assembly (Fig 7A). First, we evaluated Spike incorporation into viral particles in a single replication cycle, using a replication-incompetent (VSV-G gene was replaced with the GFP gene), vesicular stomatitis virus (VSV) model (VSVΔG-GFP, or simply VSV hereafter) that has been widely used for SARS-CoV-2 research [16,17]. Cells co-transfected with plasmids encoding Spike or ORF8-Strep were infected with VSV, and the supernatant containing VSV virions that incorporated Spike (referred to as S-VSV hereafter) was evaluated by immunoblot analysis. Significantly decreased Spike signals were detected in the S-VSV particles (normalized by VSV-M (VSV membrane protein)) produced in cells co-expressing Spike and ORF8-Strep (S(+ORF8)-VSV) than the S-VSV produced in cells expressing Spike only (Fig 7B).

**Fig 7.**
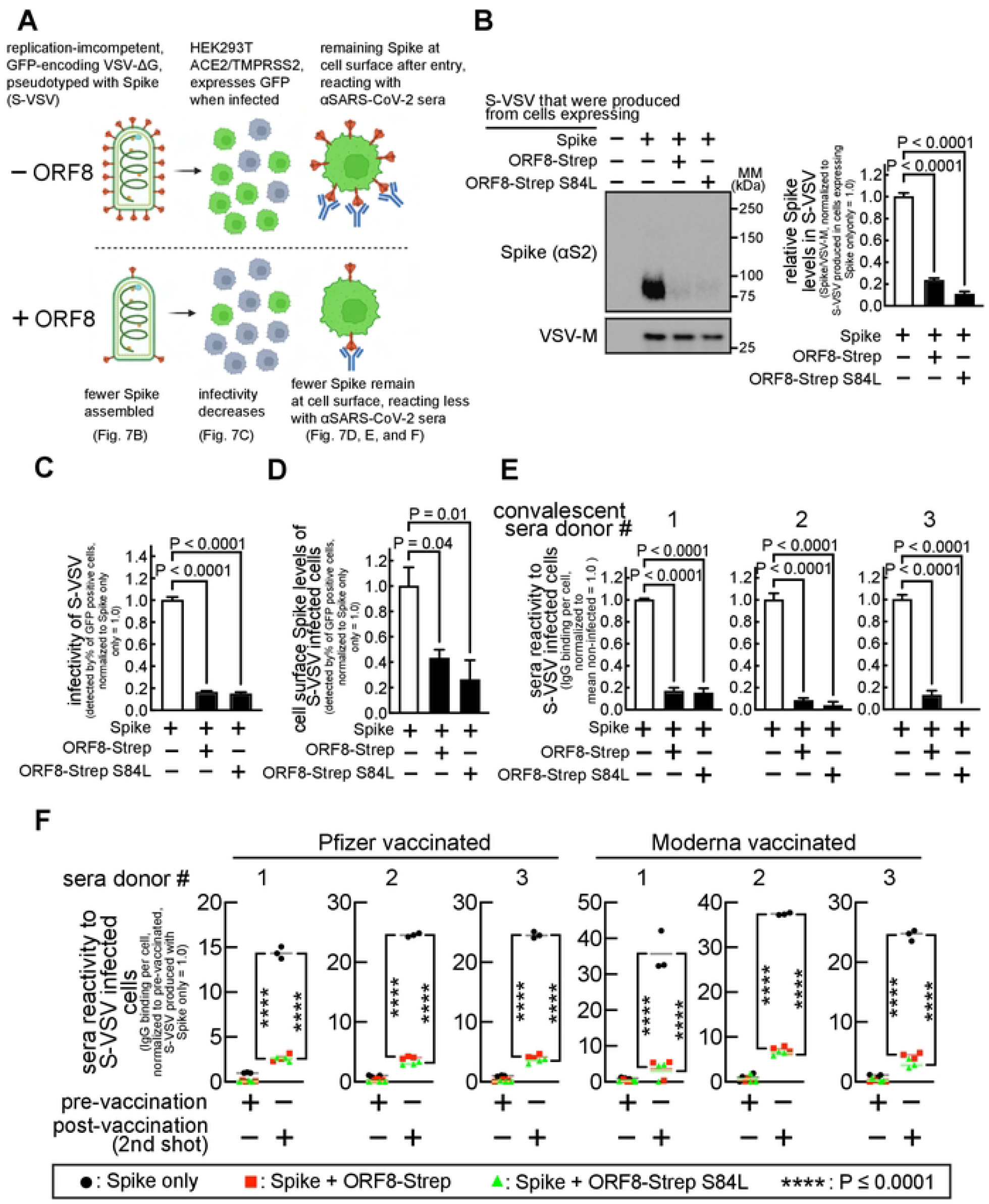
ORF8 results in decreased infectivity of Spike-pseudotyped virus (S-VSV), but limits the reactivity of anti-SARS-CoV-2 human sera towards the infected cells. **A**. Experimental workflow and summary of results. **B**. HEK293T cells co-transfected with plasmids encoding Spike or ORF8-Strep genotypes (ORF8-Strep, or ORF8-Strep S84L) were infected with replication-incompetent GFP-encoding VSV (VSVΔG-GFP), which resulted in production of Spike incorporated ΔG-GFP (S-VSV). The culture medium containing S-VSV was collected, and Spike levels in S-VSV were evaluated by immunoblot analysis with antibodies against S2 (detects both uncleaved and S2 fragment) and VSV-M (loading control, a VSV structural protein). **C-F**. HEK293T cells stably expressing ACE2ΛΓMPRSS2 were infected with S-VSV produced in the absence or presence of ORF8-Strep or ORF8-Strep S84L. **C**. The infectivity of S-VSVs were evaluated by flow cytometry analysis for the percentage of GFP expressing cells. The cells were further incubated with antibodies against S2 (detects both uncleaved and S2 fragment) (**D**), or anti-SARS-CoV-2 human sera collected from three COVID-19 convalescent (**E**), or six vaccinated (**F**) (three Pfizer and three Modema, before 1st shot (pre-vaccination) and after 2nd shot (post-vaccination)) donors. The levels of cell-surface Spike in S-VSV infected cells (**D**) and the reactivity of anti-SARS-CoV-2 human sera (**E** and **F**) towards S-VSV-infected cells were evaluated flow cytometry in S-VSV-infected (GFP-positive), but yet viable (LIVE/DEAD-negative) cells. The data represent or are combined from three independent experiments and are presented as mean ± s.d. Statistical significance was analyzed using one-way (**B–E**; Dunnett’s) or two-way (**F**; Tukey’s) ANOVA.

Next, we evaluated the infectivity of the S-VSV virions, which can be assessed by the measuring the percentage of GFP-positive cells after infection (S-VSV encodes GFP). HEK293T A/T cells incubated with the supernatant samples that contain the equal levels of S-VSV particles (confirmed by VSV-M levels) for 16 h (infection causes no cell death within this time frame) were harvested and evaluated for the percentage of the sub-populations of infected (GFP-positive) cells. We observed a significantly lower infectious unit (IU) (< 90 % decrease) in cells incubated with S(+ORF8)-VSV than cells incubated with S-VSV (Fig 7C), indicating a lower infectivity of S(+ORF8)-VSV.

Fully infectious SARS-CoV-2 particles harbor up to several dozens of Spike molecules [18]. Theoretically, only a single Spike trimer is required for cell entry [8], and we speculated that the other unreacted Spike molecules upon cellular entry remain at the cell surface. HEK293T A/T cells infected with S-VSV or S(+ORF8)-VSV (GFP-positive) in previous studies were harvested and incubated with an antibody against Spike S2. Cells were further incubated with a fluorophore-conjugated secondary antibody and a LIVE/DEAD viability dye, followed by flow cytometry analysis for the cell-surface Spike levels. Cell-surface Spike signals were easily detected in cells infected with S-VSV (Fig 7D), but reduced in cells infected with S(+ORF8)-VSV (Fig 7D). These results indicated that virus-derived cell-surface Spike is present upon infection and was lower with the viruses produced in the presence of ORF8.

Next, we examined the reaction of anti-SARS-CoV-2 sera with infected cells presenting virus-derived cell-surface Spike. The same experiment with anti-SARS-CoV-2 sera (Fig 6) (instead of anti-Spike S2 antibody) showed significantly lower (< 90%) human IgG signals in cells infected with S-VSV than cells infected with S(+ORF8)-VSV (Fig 7E: convalescent) (Fig 7F: vaccinated). These results indicated that anti-SARS-CoV-2 human sera react to the S-VSV-infected cells through virus-derived cell-surface Spike, and that the reaction was limited in cells infected with S(+ORF8)-VSV. These studies support the model that cell entry of virions produced in the presence of ORF8 leaves less cell-surface Spike, limiting reaction of anti-SARS-CoV-2 sera to infected cells.

## Discussion

The unprecedented infectivity and transmissibility of SARS-CoV-2 resulted in over 6 million deaths, in comparison to hundreds caused by SARS-CoV or Middle east respiratory syndrome. This difference suggests that SARS-CoV-2 has unique virulence mechanisms. Since ORF8 is the SARS-CoV-2 gene that is the least homologous to other coronaviruses [7], we determined if those mechanisms are mediated by ORF8 and found that ORF8 controls Spike antigen levels in virions and infected cells. Specifically, ORF8 limits production and maturation of Spike by inhibiting protein synthesis and retaining Spike at the ER. Furthermore, limited Spike levels in virions or infected cells restrict recognition by anti-SARS-CoV-2 antibodies in convalescent or vaccinated individuals, revealing a unique SARS-CoV-2 mechanism that can help evade or delay host sensing of infection.

VOCs largely emerged from rapid accumulation of pro-viral mutations, a common trait of RNA-genomic viruses [19]. Interestingly, the amino acid sequence of ORF8 is exceptionally conserved in the VOCs [9], and an ORF8-deficient variant (Δ382) from the early pandemic existed only transiently [20]. Several studies investigated the possibility that ORF8 has an indispensable pro-viral role in SARS-CoV-2 infection, but reported otherwise. The Δ382 strain replicates faster *in vitro* [20], but there is no significant change in the transcriptome of lung organoids infected with Δ382 [21]. ORF8 inhibits production of a viral component [22]. Consistently, we found that ORF8 restricts Spike incorporation into viral particles (Fig 7B), and in turn, the virions were less infectious (Fig 7C). However, our studies also revealed the ancestorial ORF8 and VOC-derived ORF8 limit reactivity of anti-SARS-CoV-2 human sera to infected cells. Therefore, our studies represent a SARS-CoV-2 strategy to control Spike antigen levels, retained through the course of evolution.

Limiting the capacity for host protein synthesis is a common viral strategy [23], hijacking building blocks and energy for synthesis of viral proteins and crippling cellular immune responses by blocking biosynthesis of immunity signaling factors [23]. Inhibition of host protein synthesis was consistently reported in SARS-CoV-2 infection [24], although the detailed molecular mechanism remains unexplored. Our studies revealed that ORF8 is the corresponding SARS-CoV-2 factor and sufficient to induce inhibition of host protein synthesis (> 90%) (Fig 4F, 4G) without requiring other SARS-CoV-2 factors. Since total Spike levels did not decrease with the non-ER ORF8 mutant (Fig 3A), we speculate that protein synthesis inhibition is linked to ORF8 cellular actions at the ER.

Cell-surface Spike and syncytia formation are evident in COVID-19 patients [14] and may allow viral spread in a manner obviating the full viral replication cycle. However, syncytia formation in SARS-CoV-2 infection induces innate immune responses through the cGAS-STING pathway [25]. Our finding that ORF8 limits syncytia formation suggests that ORF8 limits the syncytia-mediated viral spread, but prevents syncytia-dependent induction of innate immune responses. That is consistent with our model that ORF8 creates a more secured viral replication environment at the expense of infectivity.

In addition to triggering Fc receptor functions, cell-surface Spike antigens may contribute to activation of immune cells [26]. In particular, natural killer (NK) cells, key players of host immune responses to SARS-CoV-2 infection [27], are activated by integration of various activating and inhibitory receptor signals [26], including IgG Fc-specific CD16 receptor that activate NK cells upon interaction with Spike-bound IgG molecules [3,26]. On the other hand, NK cell activation can be regulated by the levels of cell-surface MHC-I molecules of infected cells. Suppressing MHC-I presentation of viral antigens, as demonstrated with SARS-CoV-2 ORF8 [28], is a powerful immune evasion strategy of several viruses [29] that, however, is programmed to be counteracted through activation of NK cells [30]. Specifically, the MHC-I-specific, killer-cell immunoglobulin-like receptor (KIR) relays inhibitory signals upon interaction with MHC-I [30]. Therefore, lack of cell-surface MHC-I molecules restricts the KIR inhibitory inputs, unleashing NK cells to activation. We speculate that limited Spike antigen levels suppress the CD16 activating signals that can, in part, counter-balance against the KIR-dependent activation, therefore, maintaining the NK cell-activating stimulations below the threshold.

Our studies revealed that ORF8 controls Spike antigen levels by inhibiting global protein synthesis and interfering with ER-Golgi process. We speculate that these cellular actions can be extended to a large number of host proteins. Especially, major immune signaling factors and receptors that are translocated into the ER for processing [31]. Therefore, ORF8 may interrupt cellular communications regulating host immune responses. In addition, while ORF8 inhibition of global protein synthesis could limit production of immune factors, and it may also reserve cellular resources for viral production. Lastly, ORF8 cellular actions reduce Spike levels in virions and infected cells, limiting cell-surface Spike antigen levels at a moment as early as viral cell entry and throughout the viral replication cycle. We speculate that this can help infected cells evade antibody-mediated phagocytosis and cytotoxicity actions for extended viral production.

In summary, our studies suggest a new SARS-CoV-2 model limiting antibody-mediated immunity. We highlight our finding that limiting levels of Spike, a key viral factor, could be pro-viral, which had been previously explored but not experimentally demonstrated [22]. Our unexpected finding of the ORF8 inhibition of the global host protein synthesis suggests additional pro-viral roles of ORF8. Future studies are required to characterize the mechanism underlying protein synthesis inhibition and ORF8’s effects on biosynthesis of host factors and metabolism. Lastly, our speculative model that ORF8 promotes immune evasion could be further explored in animal model. These future studies may lead to new therapeutics to neutralize the pro-viral ORF8 effect, which can complement ongoing countermeasures against the VOCs and help prevent re-infections or breakthrough infections.

## Materials and Methods

### Computational prediction of ORF8 subcellular localization

The whole ORF8 amino acid sequence (WA1/2020) [7] was analyzed using Protter (ETH, Zürich).

### Plasmid source and construction

Several plasmids were a kind gift from Nevan Krogan [7] (ORF8-Strep (Addgene #: 141390), Spike-Strep, eGFP-Strep (Addgene #: 141395)), Hyeran Choe [12] (Spike-Flag (Addgene #: 156420), Spike-Flag FKO (Addgene #: 159364), Flag-Spike-Flag (Addgene #: 156418), and David Nemanzee [32] (SARS-CoV Spike ΔC28 (Addgene #: 170447), Spike ΔC18 (Addgene #: 170442)). ORF8-Flag was constructed by replacing the double-Strep tags of ORF8-Strep with a nucleotide sequence (GACTATAAAGATGATGATGATAAA) encoding the Flag epitope (DYKDDDDK). SARS-CoV ORF8-Strep plasmids (ORF8ab-Strep, ORF8a-Strep, ORF8b-Strep) were constructed by replacing the ORF8 of ORF8-Strep with the corresponding genomic nucleotide sequence originated from GZ02 (ORF8ab) or BJ01(ORF8a, ORF8b). ORF8-Strep S84L was constructed by replacing TCC with CTG at S84 of ORF8-Strep. The non-tagged Spike was constructed by introducing the C-terminal cytoplasmic tail (C18) to Spike ΔC18. The non-tagged ORF8 was constructed by deleting the double Strep tags from ORF8-Strep. The GFP-Flag was constructed by replacing the ORF8-Strep of ORF8-Strep with the GFP-Flag open reading frame sequence (Sino Biological). ORF8-Flag I9P was constructed by replacing ATT with CCT at I9 of ORF8-Flag. The ORF8-Flag Δ1-17 was constructed by eliminating the first 17 N-terminal amino acids from ORF8-Flag. The fluorescence transfection reporter plasmids were constructed by replacing the ORF encoding Puro^R^ in the ORF8-Strep derived plasmids with a nucleotide sequence encoding eGFP or mCherry (SnapGene).

### Mammalian cell lines and culture condition

Human lung epithelia–derived A549 (ATCC, CCL-185) or human embryonic kidney-derived HEK293T (ATCC, CRL-3216) cells were maintained by incubating in Dulbecco’s Modification of Eagle’s Medium (DMEM) supplemented with 10% fetal bovine serum (Sigma-Aldrich) or Serum Plus II (Sigma-Aldrich), 1% penicillin/streptomycin (Sigma-Aldrich), in a humidified environment at 37 °C with 5% CO_2_. Cells were detached by incubating with trypsin-EDTA (0.05%) (Thermo Fisher) and seeded in well plates at an appropriate cell density not exceeding 90%. When firm cellular attachment is required with HEK293T cells, plates were pre-coated with rat-tail purified collagen (Gibco) as described by the manufacturer.

### Transfection for ectopic gene expression

Cellular transfection followed a standard forward-transfection method, which was validated in our studies to be experimentally similar with the reverse-transfection method. Briefly, transfection mixtures were prepared by mixing plasmids (1 μg total) with 1 μL of P3000 reagent (Thermo Fisher), and then with 1 μL of Lipofectamine 3000 reagent (all pre-diluted in Opti-MEM, (Thermo Fisher)) per well in 24-well plates. After a 10-min incubation at room temperature, the mixture was added to cell suspensions while seeding onto well plates. For co-transfection, the ORF8-encoding plasmids and the Spike-encoding plasmids were mixed at a ratio of 1:1 or 4:1 (for Spike-Flag derivatives, to tune down the expression level to other Spike constructs), which is in line with the studies that demonstrated higher ORF8 expression levels than Spike expression levels within SARS-CoV-2-infected cells [33].

### Fluorescence microscopy analysis

Mammalian cells that were seeded onto eight-well chamber slides (Thermo Fisher, Nunc LabTek II CC2) were fixed in PBS-buffered 4% paraformaldehyde (Electron Microscopy Sciences) at room temperature for 15 min, and then permeabilized in the blocking buffer (1% BSA and 0.1% Triton X-100 in PBS) at room temperature. After 10 min, the cells were washed twice with the blocking buffer and incubated at 4 °C with primary antibodies (mouse anti-Strep, Qiagen, Cat #: 34850, 1: 150 dilution) (rabbit anti-PDI, Cell Signaling, Cat #: 3501, 1: 200 dilution) (rabbit anti-Flag, Cell Signaling, Cat #: 14793, 1:250 dilution) (mouse anti-PDI, Thermo Fisher, Cat #: MA3-019, 1:200 dilution) (mouse anti-Spike S2, Thermo Fisher, Cat #: MA5-35946, 1: 500 dilution) (rabbit anti-TGN46, Proteintech, Cat #: 13573-1-AP, 1:200 dilution). After overnight incubation, the cells were washed three times with the blocking buffer and incubated at room temperature with fluorophore-conjugated secondary antibodies (goat anti-mouse IgG Alexa 488, Thermo Fisher, Cat #: A11001, 1:500 dilution) (goat anti-rabbit IgG Alexa 555, Thermo Fisher, Cat #: A21428, 1:500 dilution) with counterstaining dyes (DAPI (Sigma): 100 ng/mL, CytoPainter Phalloidin-iFluor 647 (Abcam): 1:1000 dilution). After 30 min, the cells were washed three times with the blocking buffer and mounted using Prolong Glass Antifade (Thermo Fisher). The slides were imaged using a fluorescence confocal microscope (Carl Zeiss, LSM700) with a 63 X or 40X objective (Carl Zeiss) and analyzed using ZEN Black edition (ver. 2.3) Software.

### Subcellular fractionation

Cells were subcellularly fractionated using the ER isolation kit (Sigma, ER0100) as instructed by the manufacturer’s protocol. Briefly, cells plated on two 15-cm plates were suspended in 1 X hypotonic extraction buffer, and incubated at 4 °C for swelling. After 20 min, the cells were centrifuged at 600 x g for 5 min and resuspended in 1 X isotonic extraction buffer. The cells were mechanically homogenized using a 7-mL Dounce homogenizer (10 strokes), and the lysate was centrifuged at 1,000 x g 10 min at 4 °C for removal of nuclear fractions. The supernatants were further centrifuged at 12,000 x g for 15 min at 4 °C, resulting in mitochondria-enriched pellet (washed two times with PBS before analysis). For isolation of the ER, the supernatant was ultracentrifuged at 100,000 x g at 4 °C for 60 min, and the ER-enriched pellet were resuspended in 100 μL of isotonic extraction buffer (ER fraction), which was analyzed by immunoblot, or further incubated in the presence of freshly prepared 0.035 or 0.2% digitonin (Sigma) for 45 min at 4 °C for evaluation of the differential solubility.

### Immunoblot analysis

Cell lysates were prepared by directly lysing monolayers of cells with western blot (WB) lysis buffer (2% SDS; 50 mM Tris, pH 6.8; 0.1% bromophenol blue; 10% glycerol; 10% β-mercaptoethanol (β-ME), all purchased from Sigma-Aldrich) or non-reducing WB lysis buffer (20 mM N-ethylmaleimide (NEM); 2% SDS; 50 mM Tris, pH 6.8; 0.1% bromophenol blue; 10% glycerol; all purchased from Sigma-Aldrich). After 10 min, the lysates were heat-denatured by incubating at 95 °C for 10 min. The proteins in the lysates were separated by SDS-polyacrylamide gel electrophoresis (SDS-PAGE) electrophoresis using gradient (4–20 %) PAGE gels (Bio-Rad, Mini-PROTEAN TGX), with a molecular mass marker (Bio-Rad) (Precision Plus Protein, Kaleidoscope, Cat #: 1610375). The proteins were electro-transferred to a polyvinylidene fluoride membrane (Millipore) using Turbo-Blot Turbo transfer system (settings: mixed MW) (Bio-Rad). After transfer, the blot was incubated at 4 °C with primary antibodies (rabbit anti-Flag, Cell Signaling, Cat #: 14793, 1:3,000 dilution) (rabbit anti-Calnexin, Cell Signaling, Cat #: 4691, 1:2,000 dilution) (rabbit anti-COX4, Cell Signaling, Cat #: 4850, 1:2,000 dilution) (rabbit anti-β-actin, Cell Signaling, Cat #: 5057, 1:5,000 dilution) (rabbit anti-Calreticulin, Cell Signaling, Cat #: 12238, 1:2,000 dilution) (mouse anti-Spike S2 or SARS-CoV Spike, Thermo Fisher, Cat #: MA5-35946, 1:2,000 dilution) (rabbit anti-anti-Spike S2, Cell Signaling, Cat #: 27620, 1:2,000 dilution) (rabbit anti-Spike S1, Cell Signaling, Cat #: 99423, 1:2,000 dilution) (mouse anti-Strep, Qiagen, Cat #: 34850, 1:2,000 dilution) (rabbit anti-ORF8, GeneTex, Cat #: GTX135591, 1:1,000 dilution) (mouse anti-VSV-M, Kerafast, Cat #: EB0011, 1:100,000 dilution), prepared in WB blocking buffer (5% skim milk (Bio-Rad) in Tris-buffered (pH 7.4) saline supplemented with 0.1% Tween 20 (Sigma-Aldrich) (TBS-T)). After overnight incubation, the blot was washed with gentle shaking with TBS-T twice (3 min each), and then incubated at room temperature with HRP-conjugated secondary antibodies ((goat anti-rabbit HRP conjugated, Cell Signaling, Cat #: 7074, 1:5,000 dilution) (goat anti-mouse HRP conjugated, Cell Signaling, Cat #: 7076, 1:5,000 dilution) prepared in the WB blocking buffer. After 1 h, the blot was washed with TBS-T with gentle shaking in TBS-T five times (5 min each). The proteins are visualized by using luminescence HRP substrates (Thermo Fisher) (SuperSignal West, Pico and Femto mixed at 1:1 ratio), which were captured using ChemiDoc XRS+ imaging system (Bio-Rad), imaged and quantified using ImageLab (Bio-Rad) (ver. 6.1.0.).

### Immunoprecipitation of Flag-tagged proteins

Monolayers of mammalian cells were briefly washed with phosphate-buffered saline (PBS) (Corning) and then lysed by incubating at 4 °C with IP lysis buffer (20 mM NEM, 1% NP-40 alternative (Millipore) in IP buffer base (50mM Tris-HCl (pH 7.4), 150 mM NaCl), supplemented with 1 X Halt^™^ Protease and Phosphatase Inhibitor Cocktail (Thermo Fisher). After 5 min, the lysates were collected and the cell debris were removed by centrifuging at 300 x g for 3 min. The clear supernatants were collected and prepared for immunoblot by mixing with the equivalent volume of 2 X WB lysis buffer (or 2 X non-reducing WB lysis buffer), or, further incubated with anti-Flag magnetic beads (Sigma-Aldrich, Cat #: M8823) at room temperature. For pre-treatment, the lysates were incubated with 2 % SDS (for denaturation) at 95 °C for 5 min, in the absence or presence of 0.2 % β-ME (for reduction). After 1 h, the mixture was separated using a magnetic separator and the beads were washed with IP wash buffer (0.05 % NP-40 substitute in IP buffer base) three times, and incubated in the WB lysis buffer (or the non-reducing WB lysis buffer) at 95 °C. After 5 min, the proteins eluded from the beads were analyzed by immunoblot.

### Blocking furin cleavage of Spike

HEK293T cell suspension were seeded onto well plates in the presence of 50 μM CMK (Tocris). After 18 h upon confirming no signs of morphological change, the cells were lysed for immunoblot analysis.

### Measurement of mRNA levels

Total cellular RNA samples were prepared using a Quick-RNA Mini-Prep (ZYMO research, Cat #: R1055). RNAs were then reverse-transcribed into cDNAs using iScript^™^ Reverse Transcription Supermix (BioRad, Cat #: 1708841). The relative abundance of Spike transcripts was quantified by quantitative real-time PCR (qPCR) using CFX384 machine (BioRad) with a fluorescence reporter (Thermo Fisher, Maxima SYBR Green/ROX, Cat #: K0223) and a pair of Spike-Flag specific primers (forward: GGTGCTGACTGAGAGCAATAA, reverse: CACATTAGAGCCGGTTGAGTAG, designed by using PrimerQuest (IDT)), which was quantified by calculating the 2^-ΔCt^ (normalized by the relative signals corresponding to β-actin in a separate qPCR). A robust ORF8-Strep transcription was confirmed by RT-qPCR in cells transfected for co-expressing Spike-Flag and ORF8-Strep.

### Flow cytometry analysis

Mammalian cells were briefly washed with PBS and incubated with Accutase (Gibco) at 37 °C. After 3 min, detachment of cells was aided by gentle pipetting after addition of 2 times the volume of ice-cold FC buffer (1% BSA in ice-cold PBS). The cells were transferred to a V-bottomed 96-well plate and centrifuged using a bucket rotor at 150 x g at 4 °C for 1 min. The cells were resuspended in 150 μL of FC buffer by gentle pipetting. The fluorescent signals from individual cells were detected using a multichannel flow cytometer (Cytek, Aurora), and measured using SpectroFlo (Cytek, version 3.0.1). The raw flow cytometry data was rendered using FlowJo (ver. 10.8.1.) (BD) and GraphPad Prism (ver. 9.0) (GraphPad Software).

### Measurement of protein synthesis activity

Cells were pre-incubated in the absence or presence of 10 μg/mL of puromycin (Sigma). After 5 min, the culture medium was removed, and the cells were incubated in DMEM lacking glutamine, methionine, and cysteine (Thermo Fisher), supplemented with 4 mM L-glutamine, 200 μM L-cysteine, and 50 μM HPG (Jena Bioscience) in the absence or presence of 10 μg/mL of puromycin at 37 °C with 5% CO_2_ in a humidified environment. After 30 min, the cells were collected as described under the “Flow cytometry analysis”. The cells were fixed by incubating in PBS-buffered 4% paraformaldehyde (Electron Microscopy Sciences) at room temperature for 15 min, and then permeabilized in 1 X Saponin-based permeabilization buffer (Thermo Fisher). After 10 min, the cells were centrifuged at 150 x g for 1 min, and then resuspended in the labeling buffer (prepared using components in the Click-iT Plus EdU Alexa Fluor 594 imaging kit (Thermo Fisher)). After 30 min, the cells were centrifuged at 150 x g for 1 min and resuspended in 150 μL of FC buffer for flow cytometry analysis. The subpopulation of cells that are singular (by gating FCS-A/SSC-A, then FCS-A/FCS-W), and transfection-positive (eGFP positive) were evaluated for the fluorescence signals corresponding to the cellular incorporated HPG.

### Evaluation of syncytia formation

Suspension of HEK293T A/T cells were seeded onto eight-well chamber slides with a transfection mixture. After 16 h, the cells were prepared and evaluated as described under the “Fluorescence microscopy analysis”.

### Measurement of the cell-surface Spike levels or reactivity of anti-SARS-CoV-2 human sera

Cell pellets in a V-bottomed 96 plate, prepared as described under the “Flow cytometry analysis”, were resuspended in 100 μL of FC buffer containing primary antibodies (mouse anti-Spike S2, Thermo Fisher, Cat #: MA5-35946, 1:500 dilution) (mouse anti-Flag M2, Sigma, Cat #: F1804, 1: 500 dilution), or anti-SARS-CoV-2 human sera (1:100 dilution) (COVID-19 negative, RayBiotech, Cat #:CoV-VP1-S-100) (COVID-19 convalescent, Innovative Research, Cat #: ISERSCOV2P100UL) (Vaccinated, RayBiotech, Cat #: CoV-VP1-S-100, CoV-VM1-S-100) (Supplementary Table 1). After a 1-h incubation at 4 °C with occasional shaking, the cells were washed two times by centrifuging at 150 x g for 1 min and then resuspending in 100 μL of FC buffer. After washing, the cell pellets were resuspended in 100 μL of FC buffer containing LIVE/DEAD violet dye (Thermo Fisher, 1:1,000 dilution) and secondary antibodies (goat anti-mouse IgG Alexa 647 conjugated, Thermo Fisher, Cat #: A28181, 1:500 dilution) (goat anti-human IgG Fc Alexa 488 conjugated, Thermo Fisher, Cat #: H10120) (goat anti-human IgG (H + L) Alexa 647 conjugated, Thermo Fisher, Cat #: A21445, 1:500 dilution). After 30 min, the cells were washed once by centrifuging at 150 x g for 1 min and then resuspending in 150 μL of FC buffer by gentle pipetting. The samples were then analyzed by flow cytometry with a gating strategy to specifically evaluate the sub-populations of singular (by gating FCS-A/SSC-A, then FCS-A/FCS-W), viable (LIVE/DEAD staining-negative), and transfection-positive (eGFP- or mCherry-positive) cells for the fluorescence signals corresponding to cell-surface Spike or cell-surface-bound IgGs derived from the anti-SARS-CoV-2 sera.

### Measurement of relative levels of Spike translocation to cell surface

Relative cell-surface Spike levels were evaluated as described under the “Measurement of the cell-surface Spike levels or reactivity of anti-SARS-CoV-2 human sera”. Cells for evaluating total cellular Spike levels were prepared by using the intracellular fixation and permeabilization buffer set (eBioScience), followed by the same immunostaining procedure for the cell-surface Spike levels.

### Experiments using S-VSV

The workflow scheme (Fig 7A) was created with BioRender.com. HEK299T cells were incubated with VSV-G-complemented VSVΔG-GFP (G*-VSVΔG-GFP) (Kerafast, Cat #: EH1019-PM) at the infectious unit (IU) of 3. After 20 h, the supernatant was collected, and cell debris were removed by centrifuging at 300 x g for 1 min at room temperature. The clear supernatant containing S-VSV was either concentrated using 100 MWCO Amicon Ultra-centrifugal units for immunoblot analysis, or kept at −80 °C until further infection experiment. For infection, the culture medium containing S-VSV were pre-treated to neutralize any residual G*-VSVΔG-GFP by incubating with anti-VSV-G antibody (Millipore, Cat #: MABF2337, 1:1,000 dilution) for 15 min at room temperature. HEK293T A/T cells that were plated no higher than 90% density were incubated with S-VSV with the targeted IU of 0.1–0.15. After 16 h, the cells were collected and prepared as described under the “Measurement of the cell-surface Spike levels or reactivity of anti-SARS-CoV-2 human sera”. The infectivity was measured by evaluating the percentage of GFP-positive cells, which were also evaluated for the cell-surface Spike levels or reactivity of anti-SARS-CoV-2 human sera. The studies resulted in the IU (infectious unit) less than 0.12 ± 0.01 (s.d.), where, based on a normal Poisson distribution, the probability of cells infected by a single particle is at least 94.1% (by two particles = 5.7%, by three particles = 0.2%), validating a strong linear correlation of the percentage of GFP-positive cells with the infectivity of the viral particles.

### Quantification and statistical analysis

All experimental data presented in our studies are representative of, or combined from, at least three biologically independent experiments. Immunoblot bands were quantified by densitometry analysis using ImageLab. Pearson’s coefficient between Spike and Golgi was measured within the circular area immediately encompassing the Golgi area, using ZEN Black edition. A total of 30 cells per condition (10 each from experimental replicate) were randomly selected and were subject to analysis. For flow cytometry, signals from > 10,000 corresponding cells after gating were measured to calculate the mean fluorescence intensity (MFI) per cell in each experimental replicate (For S-VSV infected cells, > 1,000 corresponding cells after gating were measured). Statistical analyses were performed using GraphPad Prism, with error bars indicating standard deviations. P values were calculated using either the Students’ *t* test with paired, two-tailed distribution, or, one-way or two-way ANOVA, corrected using either the Dunnett’s or the Tukey’s test. P values smaller than 0.05 were considered statistically significant (a = 0.05)

## Acknowledgments

We thank Frank Soveg, Irene Chen and Jennifer Hayashi in Ott lab for discussion, James Hurley at UC Berkeley for intellectual insights, David Gordon at UCSF-Gladstone for the plasmid resources, and Marius Walter and Nicolas Andrews and Rebeccah Riley in Verdin lab for assisting the project. We thank scientists who participated in the QCRG COVID-19 consortium for scientific communications. We thank the morphology core and the flow cytometry core at the Buck Institute for Research on Aging for technical support. This project was financially supported by the internal research funds at the Buck Institute for Research on Aging. M.O. graciously received support from the James B. Pendleton Charitable Trust, Roddenberry Foundation, and P. and E. Taft.

## Author contributions

I-J.K. and E.V. conceived the project and designed the experiments. I-J.K., Y.L., M.M.K., and Y.Z. performed experiments. I-J.K. analyzed and rendered the data. M.O. and E.V. provided guidance and funded the project. I-J.K. and E.V. wrote the manuscript.

---

**Fig S1.**
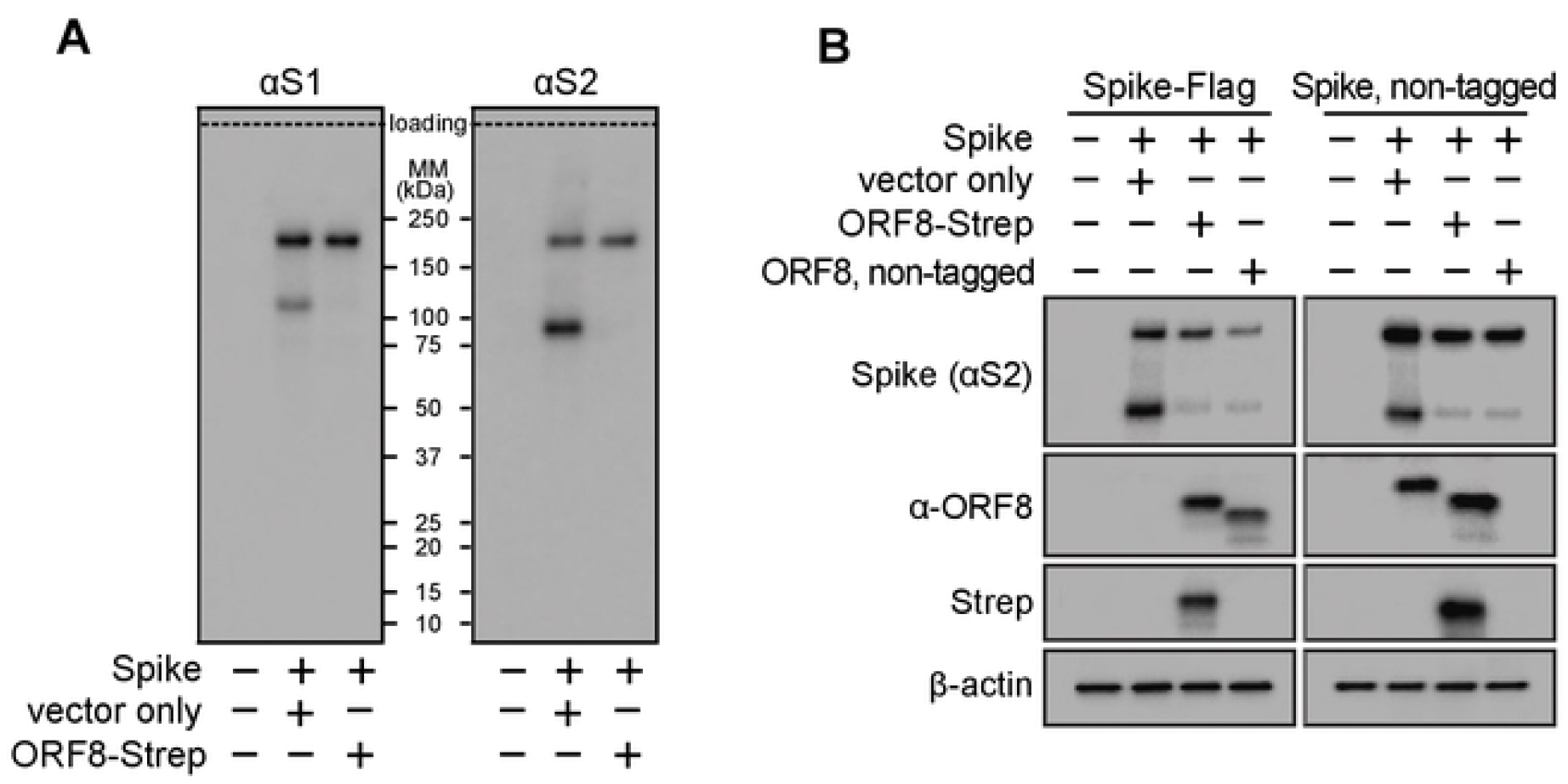
Validation of using C-terminal-tagged Spike or ORF8 constructs. **A**,**B**. HEK293T cells co-transfected with plasmids encoding Spike-Flag (**A** and **B**) or non-tagged Spike (**B**) or ORF8-Strep (**A** and **B**) or non-tagged ORF8 (**B**) were lysed and evaluated by immunoblot analysis using antibodies against S2 (detects uncleaved and S2 fragment of Spike) (**A** and **B**), S1 (detects uncleaved and S1 fragment of Spike), Strep (detects ORF8-Strep) (**B**), ORF8 (detects both ORF8-Strep and non-tagged ORF8) (**B**) and ß-actin. The data represent three independent experiments.

**Fig S2.**
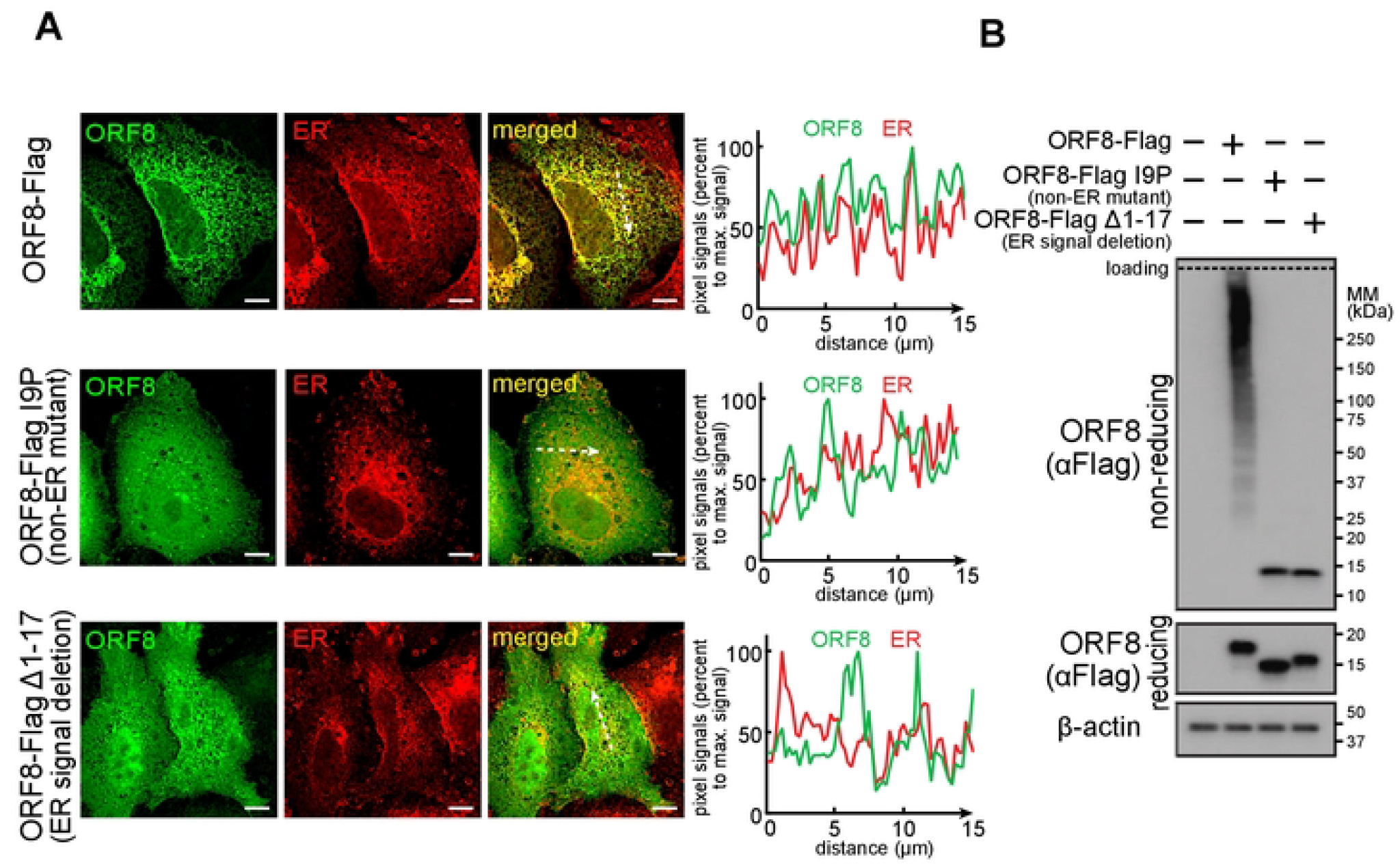
19P mutation incapacitates ORF8 translocation to ER. **A**. A549 cells transfected with a plasmid encoding ORF8-Flag, ORF8-Flag 19P (non-ER mutant), or ORF8-Flag Δ1-17 (ER signal deletion) were fixed, permeabilized, and immunostained for PDI (ER marker) and Flag (ORF8), and analyzed by fluorescence confocal microscopy imaging. White scale bars = 10 μm. The pixel intensities of ORF8 and PDI along the dashed arrow are plotted. **B**. HEK293T cells transfected with a plasmid encoding ORF8-Flag, ORF8-Flag I9P, or ORF8-Flag Δ1-17 were lysed and evaluated by immunoblot analysis using antibodies against Flag (ORF8) or ß-actin under reducing or non-reducing (protein interactions through disulfide bonds are preserved) conditions. The data represent three independent experiments.

**Table S1.** Specifications of anti-SARS-CoV-2 human sera. **A**. Source of COVID-19-negative or -positive (convalescent) sera. **B**. Source of COVID-19 vaccinated sera, three Pfizer and three Modema, collected before 1st shot (pre-vaccination) and after 2nd shot (post-vaccination).

| COVID-19 | donor # | age | sex | race | identifier # |
| --- | --- | --- | --- | --- | --- |
| negative | 1 | 32 | male | Caucasian | V901A |
|  | 2 | 48 | male | Asian | V906A |
|  | 3 | 24 | female | Caucasian | V911A |
| positive<br>(collected<br>06/17/20) | 1 | 26 | male | Caucasian | RPGG68 |
|  | 2 | 26 | female | Hispanic | TGRJLK |
|  | 3 | 26 | female | Hispanic | GT4DZ4 |

**Table S1 Specifications of anti-SARS-CoV-2 human sera. A.** Source of COVID-19–negative or –positive (convalescent) sera. **B.** Source of COVID-19 vaccinated sera. three Pfizer and three Moderna, collected before 1st shot (pre-vaccination) and after 2nd shot (post-vaccination).
| brand | donor # | age | sex | race | vaccination and identifier # |  |
| --- | --- | --- | --- | --- | --- | --- |
|  |  |  |  |  | pre- | post- |
| Pfizer | 1 | 32 | male | Caucasian | V901A | V901C44 |
|  | 2 | 48 | male | Asian | V906A | V906C11 |
|  | 3 | 24 | female | Caucasian | V911A | V911C8 |
| Moderna | 1 | 34 | male | Caucasian | V902A | V902C22 |
|  | 2 | 47 | male | Caucasian | V903A | V903C20 |
|  | 3 | 46 | female | Asian | V909A | V909C22 |

